# Reconstruction of atomistic structures from coarse-grained models for protein-DNA complexes

**DOI:** 10.1101/205062

**Authors:** Masahiro Shimizu, Shoji Takada

## Abstract

While coarse-grained (CG) simulations have widely been used to accelerate structure sampling of large biomolecular complexes, they are unavoidably less accurate and thus the reconstruction of all-atom (AA) structures and the subsequent refinement is of desire. In this study we developed an efficient method to reconstruct AA structures from sampled CG protein-DNA complex models, which attempts to model protein-DNA interface accurately. First we developed a method to reconstruct atomic details of DNA structures from a 3-site per nucleotide CG model, which uses a DNA fragment library. Next, for the protein-DNA interface, we referred to the sidechain orientations in the known structure of the target interface when available. The other parts are modeled by existing tools. We confirmed the accuracy of the protocol in various aspects including the structure deviation in the self-reproduction, the base pair reproducibility, atomic contacts at the protein-DNA interface, and feasibility of the posterior AA simulations.

## INTRODUCTION

Although numerous structures of biomolecules and their complexes have been experimentally characterized, they are mostly limited to stable structures. However, biomolecular processes are dynamic, in which biomolecules bind and dissociate each other. Transient complex structures appeared in the dynamic processes have been difficult to characterize. In this sense, molecular dynamics (MD) simulations became increasingly important to “watch” how biomolecules move and what transient complexes are formed. Yet, at the moment, the gold-standard atomistic MD simulations are limited to relatively short time scales, typically some microseconds, which is often much shorter than the biologically relevant time scales. This is especially a serious problem in the cellular nucleus processes where most proteins are highly flexible and form many transient complexes with DNA, which are keys to any molecular genetic processes^1–3^. In this context, coarse-grained (CG) MD simulation is a useful approach to study such dynamic and transient states of large protein-DNA complexes^4–11^. By reducing the resolution, the CGMD reduces the computation cost remarkably and thus enables us to explore much larger conformational space, while it is unavoidably less accurate than atomistic MD. Recently, the CGMD has been applied to various protein-DNA transient complexes, such as the full-length p53 sliding along DNA and the DnaA molecules forming the replication initiation complex at the bacterial *oriC* region^10,11^.

When we obtain novel transient structures by CGMD simulations, their accuracies are always of question. To assess the stabilities and accuracies of these structures, it must be very useful if one can reconstruct fully-atomistic structures from the low resolution models obtained by the CGMD simulations, which we call the reconstruction or the reverse-mapping. Once reconstructed, starting from the reconstructed structures we can perform posterior all-atom (AA) MD simulations, evaluate their stability, and refine the structures. Salt bridges and hydrogen bonds, for example, observed in AAMD simulations can be further confirmed by biochemical assays with mutant proteins. Furthermore, since MD simulations are inherently approximate and thus need assessment by experiments^12^. For many comparisons with experiments, the reconstructed AA structures are indispensable.

By now, some reverse-mapping methodologies for biomolecules have been developed for homology modeling, protein designs, and multi-scales simulations^13–44^. The reconstruction of protein sidechains onto given main chains is a popular example of the reverse mapping. This is usually achieved by selecting a sidechain conformation from a library of rotamers, a prepared discrete set of sidechain conformations. Many approaches have been proposed to choose suitable rotamers or conformers^31–40^. The reconstruction of a main chain of a protein from its Cα atom positions has also been studied intensively. Most methods reconstruct main chains using a fragment library taken from a set of known structures^24,30,41–44^. A geometrical method was also proposed^14^.

When the reverse mapping is discussed in the context of multi-scale simulations, often this is achieved via the two steps: The placement of atoms associated with every CG particles is followed by the energy minimization with AA force field^13–22,25–27,45^. The relative weights of the two steps vary among different methods. In one extreme, Rzepiela et al arranged atoms around each CG particle rather randomly, which is followed by the extensive simulated annealing with an AA force field^13^. On the other hand, Wassenaar et al developed a method to place higher-resolution particles with correct topology while bond lengths and/or bond angles are of low accuracy, which is followed by several rounds of structural relaxation^14^. In another example, Liu et al proposed a reverse mapping methodology based on configuration-bias Monte Carlo^23^. Notably, since a given CG particle position is usually compatible with an ensemble of AA structures, the reconstruction is not a unique process. Thus, posterior structural relaxation via AAMD is a reasonable approach to obtain a locally stable reverse-mapped structure. For CG models with relatively high resolution, atomistic details can be recovered directly by geometric calculations, where the subsequent structural relaxation is not important^28^.

In this study, we develop a new reverse-mapping method for protein-DNA complexes and test its performance. Although the method is of general nature, to make arguments unambiguous, we focus on the reconstruction of atomistic structure from a CG model of a particular resolution; one CG particle per amino acid in proteins and three CG particles per deoxyribonucleotide in DNA each representing sugar, phosphate, and base^46–48^. This resolution is taken in the general biomolecular CGMD software, CafeMol^49^, which we have been developing so that the current method can directly be applied to the output snapshots from CafeMol. Given many existing tools for the protein reverse mapping from this resolution, in this study we pay attentions to the two points; 1) the reconstruction of DNA structures and 2) the reconstruction at the interface between proteins and DNAs.

First, we constructed a DNA fragment library and fitted the most suitable nucleotide in the library to every segment in the target CG DNA structure. Both in the self-reproduction test from the crystal structures and in the test that used CGMD snapshots, we could reproduce the original structures and their interaction feature quite well. Our method reproduced the tilt of bases and Watson-Crick base pairing with high probabilities.

Accurate modeling of the protein-DNA interface is particularly important for functional studies, but, due to high atomic density at the interface, protein sidechain optimization at the interface is a rather challenging problem. When our target is the reconstruction of some transient protein-DNA complex structures for which the stable structure information is available, we can utilize that interface information as much as possible. We took the sidechain orientation in the stable (X-ray) structure only to the protein-DNA interface, while other parts are modeled with existing tools, PD2 CA2MAIN^30^ and SCWRL4^31^. To our knowledge, this hybrid reconstruction has not been examined before. In this paper, we show that the method reproduce protein-DNA interactions well. The subsequent energy minimization further improved the local structures of the reconstructed protein-DNA complexes such as bond lengths and Watson-Crick base pairings.

## METHODS

### Outline of the reconstruction method

In this work, we consider the general case where a coarse-grained (CG) model for a protein-DNA complex is given from prior CGMD simulations. Thus, the given CG model may have low-accuracy corresponding to the resolution taken in the CG simulations. Given that condition, our goal is to provide a reasonably accurate all-atom (AA) structure that is compatible with the given CG model. More specifically, we try to reconstruct an atomic structure that capture crucial inter-molecular interactions and from which standard AAMD simulations can be performed. The procedure can be divided into four parts (Fig. 1).

**Figure 1.**
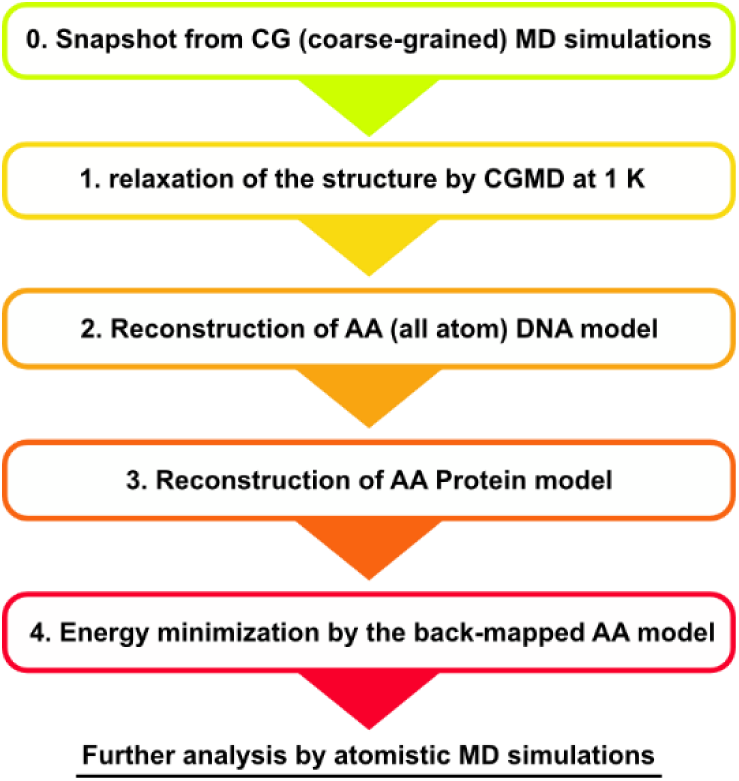
A flowchart of the reconstruction methods for protein-DNA complexes.

1. Refinement of the CG model by a CGMD simulation at low temperature (in this study, at 1 K),
2. Reconstruction of an AA DNA structure
3. Reconstruction of an AA protein structure compatible with the AA DNA structure
4. Refinement of the AA structure by the energy minimization with an AA force field.

To make the description unambiguous, we assume to use the specific CG model for proteins and DNAs. For DNAs, we use the 3SPN.2C model, in which each nucleotide is represented by three CG particles (Fig. 2A)^48^. The CG particles representing sugar, phosphate, and base are placed at the centers of mass (COM) of the corresponding groups. For proteins, we use a CG model where each amino acid is represented by one CG particle placed at the Cα atom position. We note, however, that the method can easily be generalized to other CG models.

**Figure 2.**
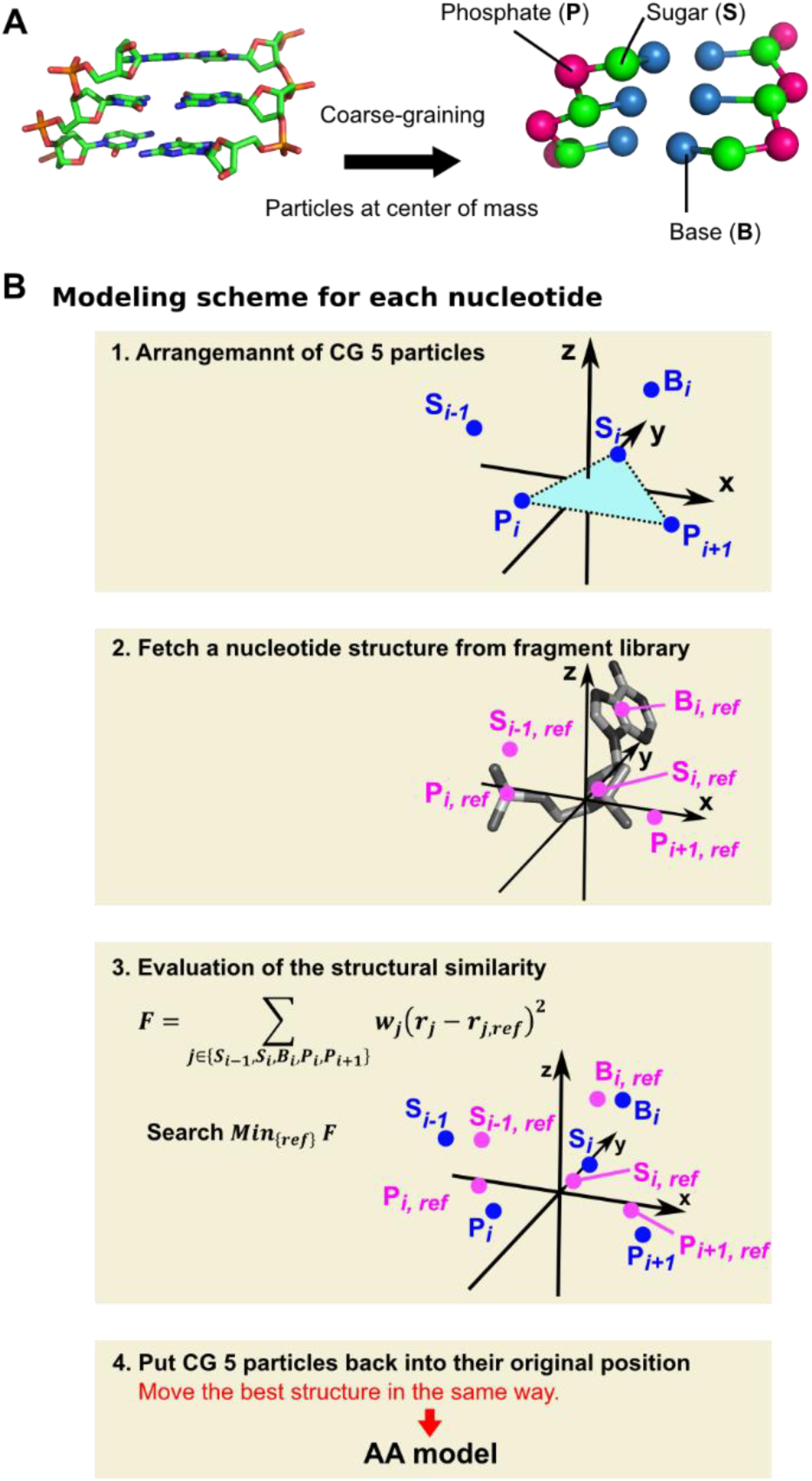
The DNA reconstruction method. A) AA and CG representations of DNA. B) The DNA reconstruction procedure.

### Preparation of the DNA fragment library

As a pre-process of the DNA reconstruction, we made a library of DNA short fragment structures.

First, we downloaded X-ray crystal structures that include DNA and that have the resolutions not larger than 2.2 Å from Protein Data Bank (PDB, the structures available on March 28^th^, 2017). We classified them into 3 groups; protein-DNA complexes with single DNA chains, protein-DNA complexes with multiple DNA chains, and the other complexes or molecules with DNA chains. For each group, we randomly chose a half set of structures and included them into the library data set (the other half set is used for the test later). For an atom with its occupancy less than 1.0 and for which more than one coordinates are given, we included the model with the highest occupancy. From each DNA structure, we extracted the structures of every deoxyribonucleotides (*i*-th nucleotide; except the 5’ and 3’-end nucleotides) in addition to their 5’-neighboring sugar groups, and 3’-neighboring phosphate and sugar groups. These correspond to 6 CG particles. The structure that corresponds to the 5 CG particles (excluding the 3’-neighboring sugar) is used for modeling non-edge nucleotides (denoted as the sugar S_*i-1*_, the phosphate P_*i*_, the sugar S_*i*_, the base B_*i*_, and the phosphate P_*i*+*1*_) (see Fig. 2B)(the nucleotide ID increases from 5’ end to the 3’ end as usual). The 3’ - neighboring sugar group is used for modeling the nucleotides at the 5’ ends. The extracted structure is called the DNA fragment in this study. We excluded the DNA fragments with missing atoms. For each DNA fragment, we calculated the coordinates of the 6 CG particles. In addition, for these 6 CG particles and all the atoms associated to these 6 CG particles, we calculated the local Cartesian coordinates **r** = (X, Y, Z) defined as the following manner. The vector pointing from Pi to P_*i*+*1*_ is parallel to the X axis. S_*i*_, P_*i*_, and P_*i*+*1*_.are placed on the XY plane. The geometric center of the S_*i*_, P_*i*_, and P_*i*+*1*_ is set as the origin (Fig. 2B). These local Cartesian coordinates of the 6 CG particles and the AA included therein are tabulated.

Collecting all the extracted fragments, we constructed the DNA fragment library, which include atom coordinates of center nucleotide, COM of base, sugar, phosphate of center nucleotide, and COM adjacent sugars, and phosphates. The PDB entries which we used for the fragment library are listed in Table S1.

As a result, 22,347 DNA fragments were prepared, in which 5,118 structures with thymine base, 5,940 structures with cytosine base, 4,980 structures with adenine base, and 6,309 structures with guanine base. We extracted a subset of fragments from the fragment library whose bases form the Watson-Crick pairing in crystal structures. The base-pairing criteria are defined below (Criteria for Watson-Crick pairing based on CG particles). We named this template group as “group A”.

### Criteria for Watson-Crick pairing based on CG particles

We defined the criteria for Watson-Click base pairing in terms of the positions of CG particles (the positions of COM of sugar and base groups for AA structures). When a pair of base particles satisfies all the following four criteria, we regarded it as forming a Watson-Crick type base pairing.

Base pairing criteria in terms of the CG representation

Adenine-thymine pair or guanine-cytosine pair which does not adjoin in DNA sequence.
The distance between two base particles is less than 6.3 Å.
We prepared representative positional relationship of 4 particles based on B-DNA structure generated by NAB, which are 2 base particles and 2 sugar particles of base-pairing deoxyribonucleotides^50,51^. The RMSD is less than 0.65 Å when corresponding 4 particles are superposed on the representative model.
When a pyrimidine base belongs to several pairs satisfying the standard above, a pair with smallest RMSD to the representative model is chosen.

In CGMD, the CG particles tend to fluctuate more largely than the corresponding positions in AA molecules. In order not to exclude base pairs which should form Watson-Crick pairing in the process of reconstruction, these are slightly lax criteria (See Fig. S1).

### Reconstruction of an AA DNA structure

Given a CG model of DNA and the DNA fragment library, we reconstruct AA structure of every deoxyribonucleotide one by one.

For every deoxyribonucleotide in the target CG DNA model, we calculate the local Cartesian coordinates of the 5 CG particles that include the three particles for the central nucleotide, the 5’ neighboring sugar, and the 3’ neighboring phosphate. Then, in the DNA fragment library, we seek the best template that minimizes the distance *F* from the target defined as

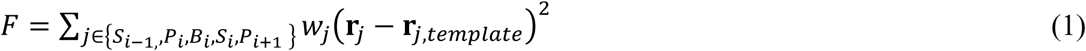

where the coefficients *w*_*j*_ weights the relative importance of 5 CG particles and are tuned later. Using the local Cartesian coordinates of the F-minimized template, we determined the position of atoms. The same procedure is repeated for all CG nucleotides.

Nucleotides at 5’ ends are modeled as follows. We calculated 6 distances, the distance between S_*i*_ and B_*i*_ (denoted as D_S*i*–B*i*_), that between S_*i*_ and P_*i*+*i*_ (D_S*i*–P*i*+*1*_), that between S_*i*_ and S_*i*+*1*_ (D_S*i*–S*i*+*1*_), that between B_*i*_ and P_*i*+*1*_ (D_B*i*–P*i*+*1*_), that between B_*i*_ and S_*i*+*1*_ (D_B*i*–S*i*+*1*_), and that between P_*i*+*1*_ and S_*i*+*1*_ (D_P*i*+*1*–S*i*+*1*_). From database, we sought the best template that minimizes the deviation *F*_*5’*_ given by,

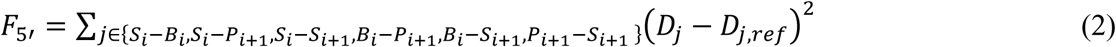

For the best template, we fitted the fragment to the CG model using S_*i*_, B_*i*_, S_*i*+*1*_, and P_*i*+*1*_. The reverse-mapped structure is the atom coordinates of the nucleotide fragment after superposition. The 3’ end nucleotide was reconstructed in the same way with *F*_*3’*_ defined in the same way as (2) using S_*i-1*_, P_*i*_, S_*i*_, and B_*i*_ positions.

Prior to calculate the *F*, *F*_*5’*_, and *F*_*3’*_, we searched Watson-Crick pairings based on the positions of CG particles and divided CG deoxyribonucleotides into 2 groups; its base particle forming base-pairing or not. For the former group, we used only the group A structures in the fragment library. For the latter group, we used all the templates in the fragment library.

### Reconstruction of an AA protein structure compatible with the AA DNA structure

We reverse mapped a sampled transient CG protein model in which each CG particle is located at the Cα position of each amino acid to obtain an AA structure. We assume here that there exists high-resolution structure information for the target protein in its stable form. For this, we take three complementary approaches. First, we superpose the high-resolution structure on the given CG model using Cα atom positions. In this procedure, we superposed the entire protein complex on CG model as a rigid body. This procedure is denoted as “Superposition”. Second, we apply existing modeling tools to reconstruct the AA details. Specifically, we modeled main chains and Cβ atoms by PD2 CA2MAIN without minimization, followed by the sidechain reconstruction by SCWRL4^30,31^. While sidechain modeling, we use the pre-reconstructed AA DNA structure as an additional steric boundary, to which hydrogens are added by gmx pdb2gmx program^52,53^. This is denoted as “PD2+SCWRL4”.

Third, we develop a hybrid approach to the case that there exists a reference known structure of the protein-DNA complex (outline depicted in Fig. 3). For the residues in the vicinity of the interface to DNA, we take the sidechains conformations from the known 3D structure and superpose them on the target CG structure. We note that when a heavy atom in an amino acid is within 7.0 Å from one of DNA atoms, we regard the corresponding amino acid as the interface to DNA. For the rest of amino acids, we simply use existing tools as above. The procedure goes in the following way. First, we reconstruct main chain AA structure by PD2 CA2MAIN without minimization. Next we superpose a known protein-DNA structure on the CG model using Cα atom positions of the entire protein complex. For each amino acid at the protein-DNA interface, we displaced in parallel the amino acid so that its Cβ atom is matched to the reconstructed backbones. Thus, the structure at the interface to DNA is composed of main chain atoms modeled by PD2 CA2MAIN and sidechain atoms by superposition followed by the parallel translation. For the rest, we reconstructed sidechains by SCWRL4. In SCWRL4, we specified reconstructed atomistic DNA and amino-acid sidechains at the interface to DNA as steric boundary condition and treated amino acids at the interface to DNA as glycine. Before applying SCWRL4, we added hydrogen atoms by gmx pdb2gmx program^52,53^.We excluded Cβ atom and Cβ atom-binding hydrogens from steric boundary condition.

**Figure 3.**
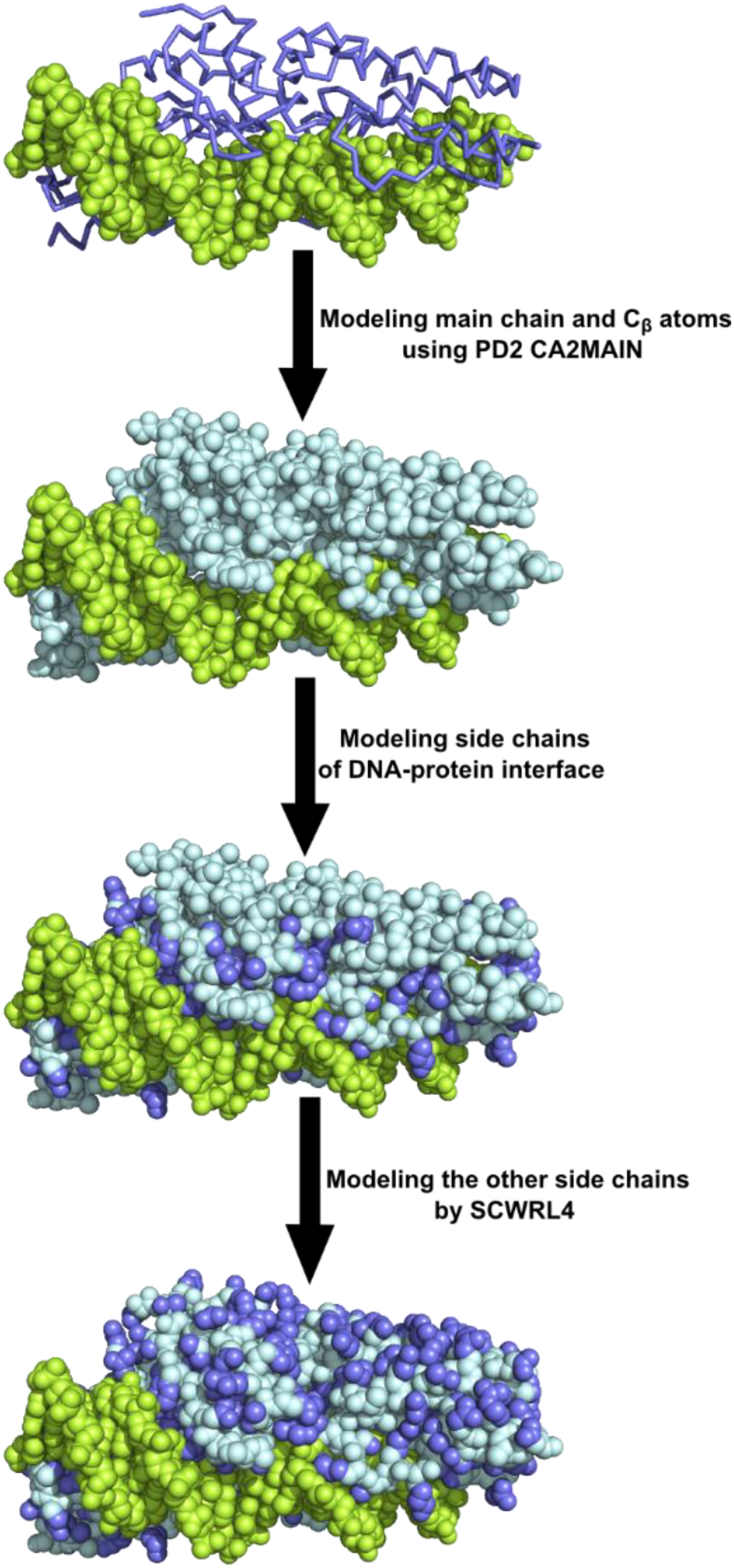
The protein reconstruction protocol. Proteins are in light cyan for main chain and Cβ atoms, and in purple for the other atoms. DNAs are in green. CG models are shown by wire and AA models are shown by spheres.

### Coarse-grained MD simulations

We used the software CafeMol 3.0 for CGMD simulations^49^. The CG model approximates proteins as a chain of amino acids, each of which is represented by one bead placed at its Cα position and DNA at a resolution of three beads per nucleotide; each representing phosphate, sugar, and base. For unambiguity, in this study, we use the AICG2+ model for proteins and 3SPN.2C model for DNA throughout although different models are applicable as well^46,48,54^. Since these models are described previously, we only briefly summarize the energy function in Supporting Information.

We performed CGMD simulations on the test set A (defined below). For each structure in the test set A, first we performed the constant temperature simulations with the Langevin dynamics at 300 K for 10^4^ MD steps. We subsequently performed the constant low-temperature simulations at 1 K for 10^4^ MD steps to refine the local structures (e.g. virtual bond length, virtual bond angle). We repeated this set of simulations 20 times changing the random seed in the Langevin dynamics.

To find an optimal condition to improve the DNA local structure, we also tested simulations at 1 K for 10^2^ MD steps and 10^3^ MD steps, as well as the simulated annealing simulations with various conditions (See Supporting Table S2, and Supporting Methods).

### Atomistic MD simulations

All atom (AA) MD simulations were performed by GROMACS 2016.3^52,53,55^. We performed energy minimization and subsequent short NVT simulation. Mainly we used AMBER99 force field with Parmbsc1^56^ for DNA, and TIP3P^57^ for water. In order to check force field dependency, we also tested the case with CHARMM27 force field^58–62^ and CHARMM TIP 3-point water model^62^. We used triclinic simulation box with periodic boundary condition. We set the box size so that the minimum distance between the DNA-protein complex and the box edge was 1.0nm. After putting DNA, the box was filled with waters, Na_+_ ions, and Cl^-^ ions. Ions are added up to salt concentration of 100mM. The particle mesh Ewald was employed for electrostatic interaction with real-space cut-off for electrostatic interaction was 1.0 nm^63^, and cut-off for Lennard-Jones potential was set to 1.0 nm. Energy minimization was conducted by steepest descent algorithm. After minimization, we performed the NVT simulation at 300K for 10 ps with the step size of 2fs. This is enough to test the stability of the modeled structures. We restrained all heavy atoms. All the covalent bond lengths in proteins and DNAs are constrained by LINCS^64^, and for waters by SETTLE^65^. For thermostat, we used the velocity-rescaling method developed by Bussi et al with coupling time of 0.1 ps^66^. In this study, we performed these operations for 900 DNA-protein complexes. The test set A contains 180 X-ray crystal structures. For each crystal structure we simulated from 5 different structures obtained by independent CGMD simulations and the subsequent reverse mapping.

### Evaluation of our modeling method

#### Dataset for testing the reconstruction method

For testing our methods, we used the following three sets of structures. The PDB entries are listed in Table S3.

**Test set A:** This is the primary test set. We prepared 180 protein-DNA complex structures. This set was used for all data except those in Fig. 4.

**Figure 4.**
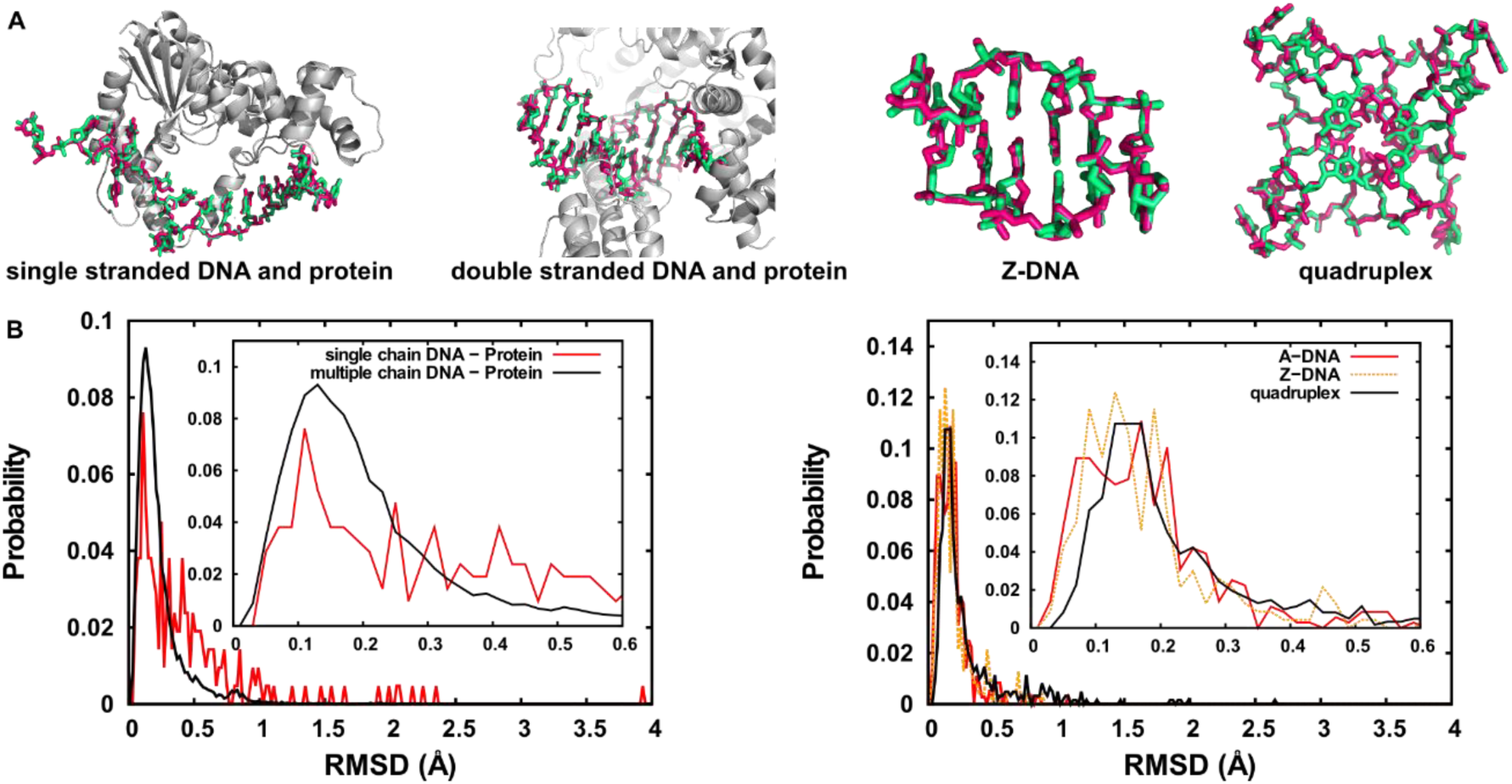
Self-reproduction test of various DNA structures. We tested single-stranded DNA in complex with proteins, multiple-chain DNA in complex with proteins, A-DNA, Z-DNA, and quadruplex DNA. A) Comparison between X-ray crystal structures (pink) and reconstructed structures (green). B) The distribution of RMSD between the original X-ray crystal structures and the self-reproduced AA structures for nucleotides that are not at the terminus in single-chain DNAs and multiple-chain DNAs (left) and in A-DNAs, Z-DNAs, and quadruplex DNAs (right).

**Test set B:** This is used to test reconstruction of non-canonical DNA in Fig. 4. We prepared 30 protein-DNA complex structures with single stranded DNA, 642 protein-DNA complex structures with multiple DNA chains, 39 A-type DNA structures, 29 Z-type DNA structures, and 22 G-quadruplex structures.

**Test set C:** This contains 25 structures, which is a subset of the test set A. This was used for determining the schedule to refine the local structure of DNA, and to check the force field dependency of energy minimization at the atomistic resolution. We chose the 25 structures from the test set A randomly excluding protein-DNA complexes that are too large, or contain too few interface contacts. The latter is because those structures tend to change interface positional relationship rather quickly, and thus they are not good for checking the global structural change during refinement CGMD run.

All the structures used are X-ray crystal structures with the resolution not larger than 2.2 Å. Structures used in the DNA fragment library were excluded. In order to perform reverse mapping and MD simulations easily, we screened the structures and edited the coordinate file. In test-set A, we excluded any structures in which DNA contained the internal deoxyribonucleotides with missing atoms or with base other than adenine, cytosine, guanine, or thymine. We also excluded any structures which contained internal incomplete amino acids or internal residues other than canonical 20 species or selenomethionine (selenomethionines were treated as methionines). If there is a missing atom in terminus nucleotide or amino acid, we used the structure after omitting the nucleotide or amino acid. In test-set B, we excluded any deoxyribonucleotides with missing atoms or with base other than adenine, cytosine, guanine, or thymine. For test set A and C, we simply used the molecules explicitly given in the pdb files, regardless of their functional units.

#### The root mean square deviation (RMSD)

As a first and quick test of the DNA reconstruction, we converted DNA X-ray crystal structures into the CG DNA model (3SPN.2C model), followed by reconstruction of the AA DNA model. To evaluate the reconstruction performance in this process, we utilized the root mean square deviation (RMSD) of each nucleotide between the initial crystal structure and the reverse-mapped structure. The RMSD was calculated without rotation or superposition of each nucleotide.

#### Base-pairing reproducibility

When a CG DNA model forms base-pairings, our reverse-mapping method should reproduce Watson-Crick pairing hydrogen bonds at the atomic resolution. We use it as a mean of evaluation of the reconstruction method, comparing base-pairings in the reverse-mapped structure with those in the CG structure. For each model, we calculated the ratios of the atomically reproduced base pairings relative to the total number of the base pairings in the CG structure. Subsequently we calculated the distribution of the ratio. For CG models, the criteria for the base pairings are shown above. When several “base-pairs” we defined share the same purine base, we referred reverse-mapped model and adopted the pair whose maximal donor-acceptor distance was minimum. In fact, these excluded base-pairs by shared purines were very rare (< 0.6 %). For reverse-mapped AA models, the criteria for the base-pairings are as follows. When the distance is less than 3.5 Å for all donor-acceptor atom pairs of a Watson-Crick pair, we regarded the two bases as forming base-pairing.

#### Joint property

Since we reverse-map DNA by each deoxyribonucleotide, the connectivity between adjacent nucleotides can be problematic. We checked if the bond lengths and bond angles between two nucleotides are close to those expected. For the test, we used the length of covalent bonds between P atom and O3’ atom, as well as the angle formed by O5’, P, and O3’ atoms.

#### Reproducibility of protein-DNA atomic interactions

We examined whether the protein-DNA interface at the initial structure was maintained after reverse mapping from a snapshot of CGMD simulations. For some atom pairs which are frequently close each other in known structures (e.g. N_ζ_ atom of lysine and OP1 atom of DNA backbone), we calculated the corresponding distances. We picked up atom pairs whose distance is less than 7.0 Å at the initial (crystal) structure and measured the distance of the atom pair. After reverse mapping from the CG MD snapshot, we calculated the distance of the same pair again. When the distance was within ±1 Å from its original value, we regarded the interaction of the atom pair was reproduced. We numerated the number of reproduced interactions in the dataset and obtained the reproduction ratio. In case that the energy minimization in the AA model could not be performed or that the subsequent NVT AAMD run stopped, we regarded as failure to reproduce all the interactions.

#### DNA helical axis

DNA helical axis of reconstructed model was calculated by Curves+^67,68^.

#### DNA reverse-mapping by another tool

For the comparison of the performance, we use a backmapping tool developed by Wassenaar et al^14^ with GROMACS 5.1.1^52,53,55^. We tested the method with PDBID 1BNA, DNA part of PDB ID 1A1F, and DNA part of PDB ID 1CDW. In order to apply the tool to the 3SPN.2C model, we changed the correspondence between CG particles and atoms. since it was unclear whether O3’ atom was assigned to phosphate particle of adjacent nucleotide, we examined the two settings; O3’ atom belongs to phosphate and O3’ atom belongs to sugar particle.

## RESULTS AND DISCUSSION

### Examination of DNA reconstruction by the self-reproduction

First, we examine the method for the reconstruction of atomic DNA structures for the target CG models that were created directly from the atomic structures (termed “self-reproduction”). In this test, we can clearly measure the performance of the method; the better the method is, the smaller the deviation between the original and the reconstructed structures is. However, we also note that, in this examination, the given CG models are limited to “high-quality” ones which are perfectly compatible with atomic structures. Importantly, this is not the case for practical applications, where CG models are created by CGMD or modeling so that the CG models are not perfectly compatible with atomic structures. Yet, we consider this self-production test is useful because it is the necessary condition for the method to be a good reconstruction tool.

First we performed the self-reproduction test for five sets of protein-DNA complexes with broad range of DNA structures (termed the test-set B); 1) 30 protein-DNA complexes with single-stranded DNA chain, 2) 642 protein-DNA complexes with multiple DNA chains (dominated by duplex DNAs), 3) 39 A-type DNAs, 4) 29 Z-type DNAs, and 5) 22 quadruplex DNAs. Here, we used the parameters *w*_*Bi*_ = 1.0 *w*_*Si-1*_ = 0.1, *w*_*Pi*_ = 0.75, *w*_*Si*_ = 1.0, and *w*_*Pi*+*1*_ = 0.1 (See Methods), which were tuned in advance. The detail of parameter tuning process is shown in Supporting Information (Supporting Results & discussion, Fig S2, S3, S4, Table S4, S5, S6).

Four representative cases of self-reproduction are depicted in Fig. 4A where the original and the reconstructed DNA structures are drawn in pink and green, respectively. These illustrate that, in most cases, our reconstruction method work well not only for B-type duplex DNA, but also for single-stranded DNA, Z-type duplex DNA, and quadruplex DNA. For each nucleotide that is not the terminus, we calculated the RMSD between the crystal structures and the reconstructed atomic models. Statistically, in every set, the RMSD distribution had its peak around 0.1-0.2 Å, suggesting that our method is applicable to most structures of DNA (Fig. 4B).

Next, using a smaller test set (the test-set A) that contain 180 X-ray crystal structures of DNA, we conducted more systematic test of reconstruction. By the same self-reproduction test as above, the distribution of RMSD (Fig. 5A) shows its peak near 0.14 Å, 70% of nucleotides had the RMSD within 0.25 Å, and 91% of them fall within 0.5 Å. Therefore our method was able to reproduce experimentally determined structures rather accurately in this test set too. By visual inspection, we found that our method can reproduce the tilt of bases even though we did not take into consideration of the base-stacking interaction or repulsive interactions between base planes, implying that the tilt of base is highly restricted in most cases.

**Figure 5.**
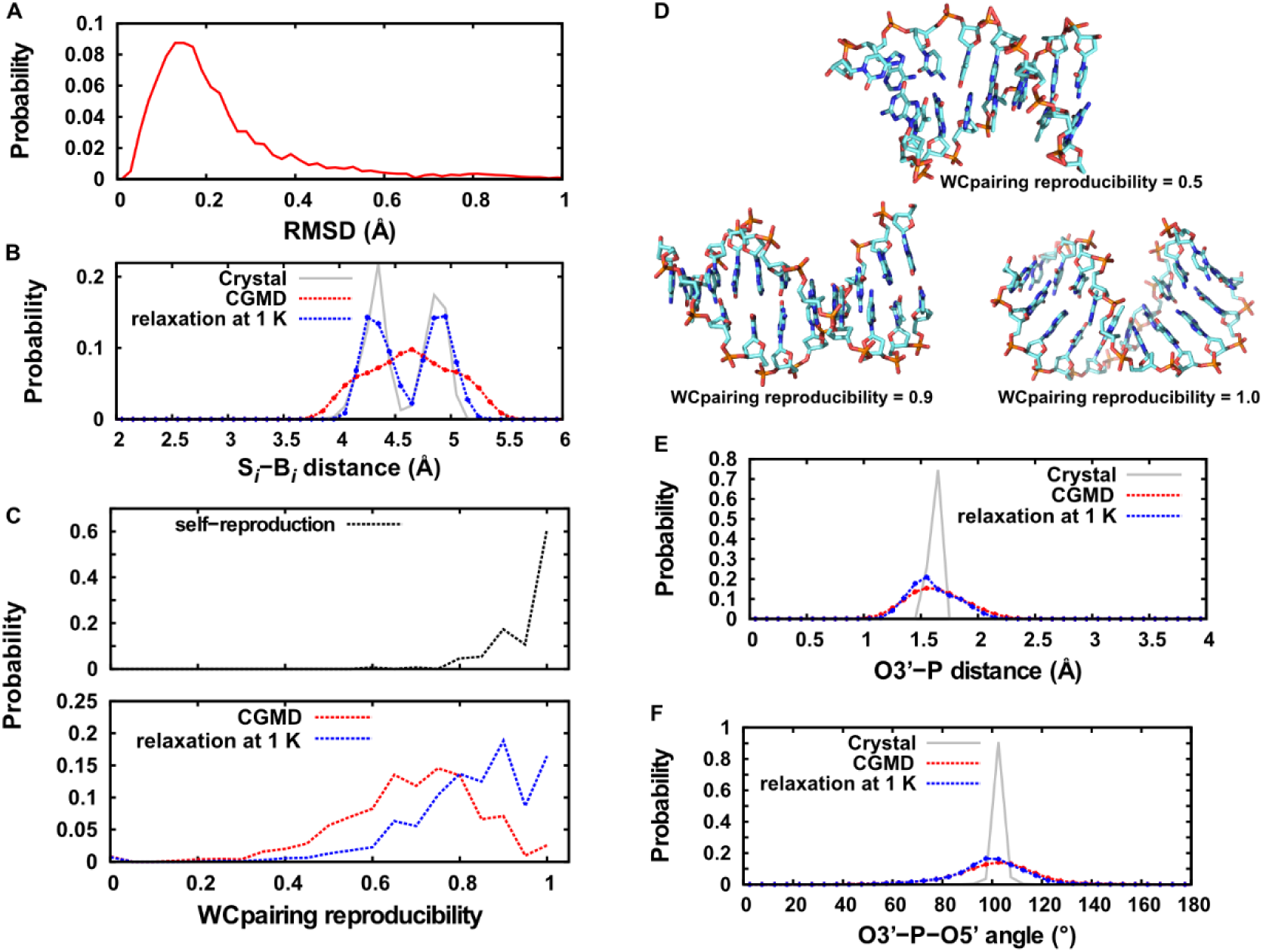
The DNA reconstruction for the test set A. A) Self-reproduction test. The RMSD distribution of inner nucleotides is shown. B) The distribution between the CG base particle and the CG sugar particle after CGMD simulations. The distribution of the distance between the center of mass (COM) of the base group and the COM of sugar of each deoxyribonucleotide in the original crystal structures are also shown in gray. C) The Watson-Crick base pairing reproducibility. D) Some examples of reconstructed DNAs with their Watson-Crick pairing reproducibility. E) The distribution of O3’-P covalent bond length. F) The distribution of O3’-P-O5’ covalent bond angle. “CGMD” means snapshots after CGMD, “relaxation at 1 K” denotes the structures after CGMD and the subsequent 10,000 MDstep-CGMD at 1 K. “Crystal” means the result for the original crystal structures. In “CGMD” and “relaxation at 1 K” cases, the error bars represent standard errors in 20 independent simulations.

As outliers in the self-reproduction of the test-set A, we found ten deoxyribonucleotides with the RMSDs larger than 2.0 Å. In these cases, bases were rotated nearly 180° along the bond connecting sugar and base groups (See Fig. S4). In these cases, the nucleotides did not form base pairing or stable base stacking in the crystal structure. Thus, the reconstruction of non-duplex DNA is at least more difficult than DNA duplexes.

Next, in the test set A, we discuss the reconstruction of the end nucleotides for which, as described in Methods, we used a different method. The RMSD for 5’ end nucleotides was modestly larger than the internal and 3’ end nucleotides (See Fig. S5A). We consider that it is easy to relax end nucleotides by posterior atomistic MD. Thus we did not consider any additional improvement.

As a quick benchmark test of computational time, we modeled 145-base pair DNA (PDBID: 3UT9) for 1000 times, which took ~140-153 seconds with Intel(R) Core(TM) i7-3770 CPU at 3.40GHz. We performed the test using single core. The time includes that for reading the fragment library and modeling 1,000 structures. Thus the computer time for the reconstruction is about 0.001 second / base pair. This is sufficiently fast, for example, to reconstruct atomic structures for all the snapshots along a CGMD trajectory.

### Examination of DNA reconstruction from CGMD simulation data

Next we tested whether our reverse mapping method is applicable to CG structures obtained from CGMD simulations. Unlike the reconstitution in the previous section, CGMD snapshots may include less accurate local structures sampled by the CGMD. As described in Methods, our reconstruction procedure begins with the short CGMD at 1 K, which is followed by the DNA reconstruction in the same way as above. We note that the CGMD at 1 K can relaxes the local structures, but does not significantly affect large-scale conformations. We searched for an optimal method to relax the local structures, including low-temperature MD and the simulated annealing with various conditions. The detail of the results is shown in Supporting Information (Supporting Results & discussion, Fig S6, S7, S8, Table S2). Finally we adopted this low temperature MD method.

For the 180 crystal structures in the test-set A, we first performed short-time constant-temperature CGMD simulations, of which the final structures are the target of our reconstruction. Subsequently, we relaxed the structure by CGMD at 1 K for 10,000 MD steps.

First we compared the CG virtual-bond distances between the base particle and the adjacent sugar particle in the initial (crystal) structures with those after the constant temperature CGMD, those after the relaxation at 1 K (Fig. 5B). In the crystal structure, there are two sharp peaks near 4.3Å and 4.9Å, which, respectively, correspond to the distances of sugar-pyrimidine base and sugar-purine base (Fig. 5B). After the CGMD, the distribution was smoothed out due to high degree of fluctuations showing only one broad peak around 4.6Å. After the relaxation 1 K-MD, the two-peak feature observed in the crystal structures recovered clearly (blue curve in Fig. 5B). Therefore, the short constant temperature MD at 1 K can improve less accurate local bond lengths appeared in the CGMD snapshots.

We also plotted the virtual bond lengths between the sugar particles and the adjacent phosphate particles in the same nucleotide. The distance also drastically improved after CGMD at 1 K (Fig. S10A). For virtual bond angles, we plotted the angle formed by phosphate, sugar, and base in the same nucleotide (Fig. S10B). Although the relaxation by the 1 K CGMD made virtually no difference, the angle after CGMD simulations showed sharper distribution than that in the crystal structures.

Next, we examined the base-pairings in the reconstructed DNA structures. First we checked Watson-Crick pairing in CG models based on our criteria (See Methods). Then we examined the reproducibility of base pairings in the reverse-mapped structures (Fig. 5C) as a test of our method. The detail definition of the reproducibility is described in Methods. When we performed the self-reproduction from the crystal structures as in the previous section, we reproduced the base pairing with very high probabilities; for 61% of DNA crystal structures in the test-set A, we reproduced all the base-pairs (Fig. 5C, black). Here, we defined the reference base-pairings using the CG model made from the crystal structure. On the other hand, for the CG models obtained from CGMD simulations, the probabilities of the base-pairing reproducibility in each target were between 0.3 and 1.0 (Fig. 5C red). Here, the reference base-pairings are defined using snapshot after CGMD at 300 K. The probability of the reproducibility markedly recovered after the relaxation MD at 1 K (Fig. 5C blue). The number of structures with the reproducibility >0.8 increased significantly, and those with the perfect reproducibility also increased. Thus, low-temperature CGMD improves the Watson-Crick pairing reproducibility. Even for the structures with reproducibility value around 0.5, the tilt of base was not so problematic in most cases (Fig. 5D). We also checked whether this Watson-Crick pairing reproducibility can be improved by larger fragment library. We performed DNA reconstruction with various sizes of the fragment library (Supporting Result & discussion, Fig. S11, Table S7). Even if we use larger fragment library, the reproducibility improved only a little. Compared to the self-reproduction test, the difference is almost negligible. The dataset dependency of the other scores is described in Supporting Result & discussion.

Finally we examined the AA local structure between adjacent nucleotides in the reconstructed AA DNA models. Notably, in the reconstruction, we superimposed each nucleotide AA structure one by one. Thus, while each intra-nucleotide structure is always accurate, the connectivity between two nucleotides is not guaranteed. We calculated the distances between O3’-P that should make covalent bonds (For connectivity between inner nucleotides in Fig. 5E, and that between 5’- or 3’ end and inner nucleotide in Fig. S5BC) and the bond angle for O5’, P, and O3’ atoms (Fig. 5F and S5DE). We found that the peaks of the distributions were well maintained for both distances and angles although the distributions became broader than those in the crystal structures. In these cases, the relaxation MD at 1 K did not improve the quality of these local structures at boundary of two nucleotides. We considered that our reverse mapped models are good enough to subsequent atomistic MD simulations. In fact, these values were drastically improved by minimization with AA force filed, as discussed below.

### Examination of the protein reconstruction at the interface to DNA

In the protein reconstruction, accurate modeling of the sidechain orientation at protein-DNA interface is of crucial importance. We consider the two cases; 1) protein-DNA complexes of which stable structures are experimentally known and from which structures were perturbed during CGMD simulations while maintaining protein-DNA interaction interface, and 2) protein-DNA complexes for which new interactions are formed in CGMD simulations. In the case 1), it is probably better to use the known structural information as much as possible. For the case 2, it should be noted that, without any experimental structure of the complex, we cannot obtain accurate sidechain orientations in CGMD simulations. The preferred sidechain orientation can only be elucidated by AAMD simulations after the reverse-mapping.

Focusing on the case 1, we decided to utilize sidechain orientations of the known structure at protein-DNA interface. To maintain the interface as much as possible and to account for significant structural change in proteins, we devised a hybrid approach where sidechain orientation of known protein-DNA complex structure is applied only to interaction interface. The rest of protein region is modeled by the existing general sidechain modeling tool, SCWRL4.

We performed the protein reverse mapping after CGMD simulation and the DNA reconstruction. Prior to test our hybrid approach, we confirmed that the distance between two amino acid beads converged to that observed experimentally. The Cα-Cα distance distribution has sharp peak at 3.8 Å except for the case of *cis* proline. By 10,000-step of constant temperature MD at 1 K, the virtual bond lengths came close to those in the initial structure of CGMD simulations (Fig. S12). After protein reverse-mapping, we performed energy minimization.

After energy minimization by an AA force field, we tested the reproducibility of interactions between protein and DNA. We regarded 10,000-step CGMD simulations at 300 K as short enough and thus assumed that protein-DNA interface was intact at the end of CGMD simulation. In this condition, we can expect high reproducibility of protein-DNA interaction when we simply superpose the protein crystal structure on CG model. Thus we regarded the “Superposition” method as a positive control. We picked up atom pair of 15 kinds and calculated the reproducibility of the atom-atom contact after reverse mapping. The detail definition of inter-atom interaction reproducibility is shown in Methods. Results are shown in Fig. 6A and Table S8, The tables contain protein interactions to the DNA bases, important for sequence-specific recognition by proteins (yellow background color), and those to the DNA backbones, important for the sequence-nonspecific feature, including protein-DNA affinity (grey background color). Our hybrid approach presented similar quality to the overall superposition method with maintenance of Cα positon at CG model. Our method successfully reproduced the sidechain orientation of crystal structure at the protein-DNA interface. At least for the 15 pairs we chose, the reproducibility of protein-DNA base was better than protein-DNA backbone interactions in our hybrid method. In many cases, the reproducibility by fully using the PD2+SCWRL4 methods was worse than the other two approaches. As contacting amino acid sidechain atoms are more distant from the main chain, the reproducibility by PD2+SCWRL4 becomes worse. For example, the reproducibility for the pair of lysine N_ζ_ atom and DNA OP1 atom is clearly low (~40%) while that of serine O_δ_ atom and OP1 atom pair is better (~60%). Probably, choice of sidechain rotamer different from known structure reflects these results. Once the main chain was modeled, the position of O_δ_ atom is nearly determined. In this condition, the method which reproduces both sidechain orientation and the position of Cα is advantageous. Therefore our hybrid approach is effective for protein-DNA complex in the case 1 situation.

**Figure 6.**
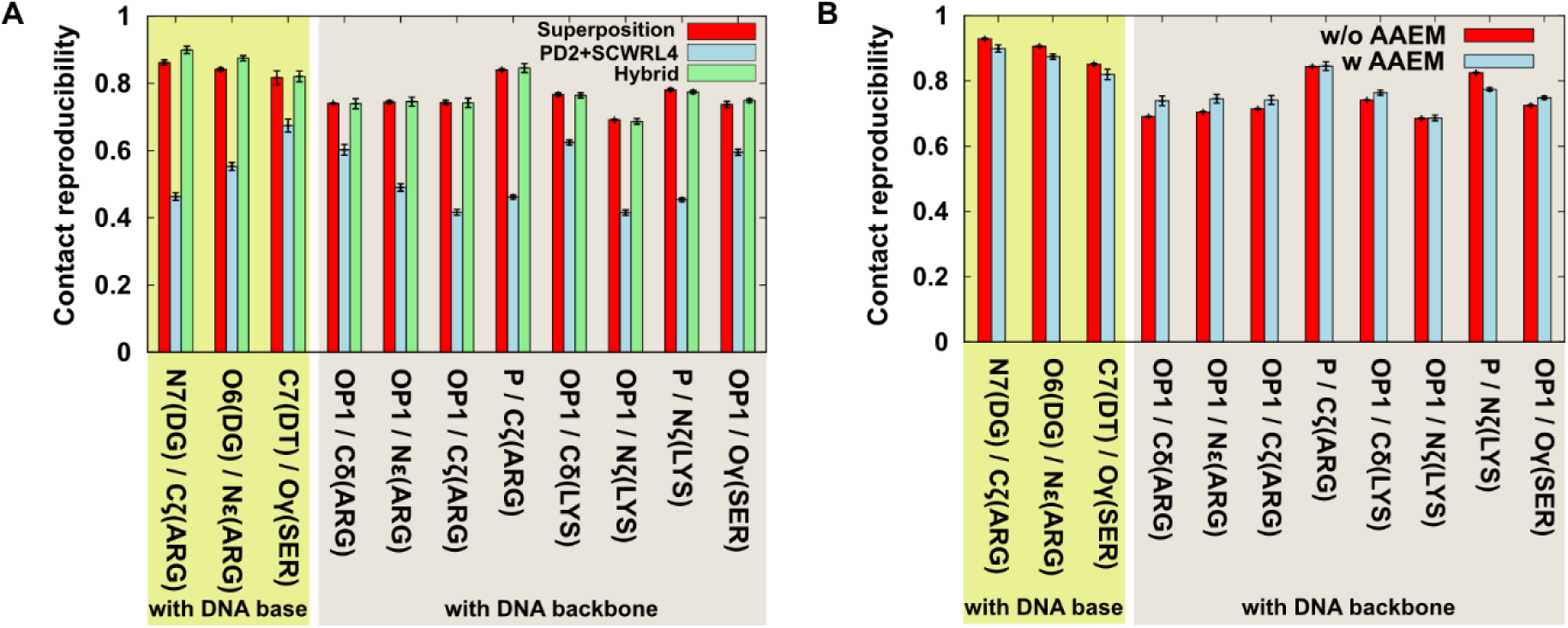
The reproducibility of atom contacts at protein-DNA interface. A) The contact reproducibility of reconstructed models with the three different protocols. The error bars represent standard errors in 5 independent simulations. “Superposition”; the high-resolution structure is superimposed on the given CG model using Ca atom positions. “PD2+SCWRL”; PD2 CA2MAIN is used to model main chains and Cβ atoms and SCWRL is used to model sidechain conformations. “Hybrid”; PD2 CA2MAIN is used to model main chains and the sidechain orientation of known protein-DNA complex structure is applied to the interface residues. “DG” represents guanine and “DT” represents thymine, respectively. The AA energy minimization was used in this calculation. B) The contact reproducibility of reconstructed models with (w AAEM) and without (w/o AAEM) the energy minimization by an AA force field. “Hybrid” approach was used here.

We also compared the reproducibility of our methods that contain the AA energy minimization (w AAEM) with that lacking the AA energy minimization step (w/o AAEM): The reproducibility was overall rather similar between “w AAEM” and “w/o AAEM” (Fig 6B, S13, Table S8, S9). The “PD2+SCWRL4” method showed relatively poor reproduction irrespective of AAEM step. Thus, when the initial structure is poor, it was not easy to improve sidechain orientations at protein-DNA interfaces merely by the AAEM. The first placement of sidechain orientation has a dominant role.

In addition we checked the AA force field dependency of DNA-protein contact reproducibility. Using the test set C, we compared the results obtained by the AMBER parmbsc1 force field with the results obtained by the CHARMM27 force field (Table S10), showing nearly no dependency at least between these two force fields.

We also checked the bond angles. Notably, the Cβ atom is the boundary of superposed structure and the main chain reconstructed by PD2 CA2MAIN, and thus the reconstruction of the correct bond angle is not trivial. Even before energy minimization, the angle distributed around that observed in the crystal structures. A subsequent energy minimization run in the AA model of protein-DNA complex made the angles close to those found in the experiment (Fig. S14). We note that our hybrid strategy is applicable also to protein-protein complexes and protein-ligand systems generally.

From the structure after the energy-minimization, we performed short constant-temperature AAMD simulations with bond length constrained by LINCS^64^. We used GROMACS 2016.3^52,53^. The MD runs completed without the error for the 95% of the cases, while 5% of the cases stopped because of unusual structure, in most case with LINCS warnings. Therefore, for majority of cases, these reverse-mapped protein-DNA complexes can be further analyzed by the posterior AAMD simulations. We note that most of the cases that stopped contained non B-form duplex DNAs, perhaps because the 3SPN.2C DNA CG model is less accurate for non B-form duplex DNA.

### Examination of the DNA structure after the energy minimization of DNA-protein complex

First we checked the base-pairing reproducibility. After the energy-minimization, the reproducibility was higher than 0.8 for almost all the complex structures and 1.0 for more than 60% structures (Fig. 7A), which is remarkably higher than that before the minimization. Surprisingly, the Watson-Crick pairing reproducibility was the same level as self-reproduction. It suggests that our method has enough performance to practical use. When we plotted the distribution of the donor-acceptor distance of the Watson-Crick pair, the distance was mainly between 2.7 to 3.2 Å, which is much narrower range than that in the self-reproduction run and that before the minimization of the reconstructed structure from CGMD runs (Fig. 7B). By taking into consideration of the AA energy, backbone structures were markedly improved: Both the O3’-P bond lengths and the O5’-P-O3’ bond angles converged to those in the crystal structure (Fig 7 C-D, S5 B-E). Thus, the posterior energy minimization with an AA force field significantly improved the quality of the protein-DNA complex AA structures.

**Figure 7.**
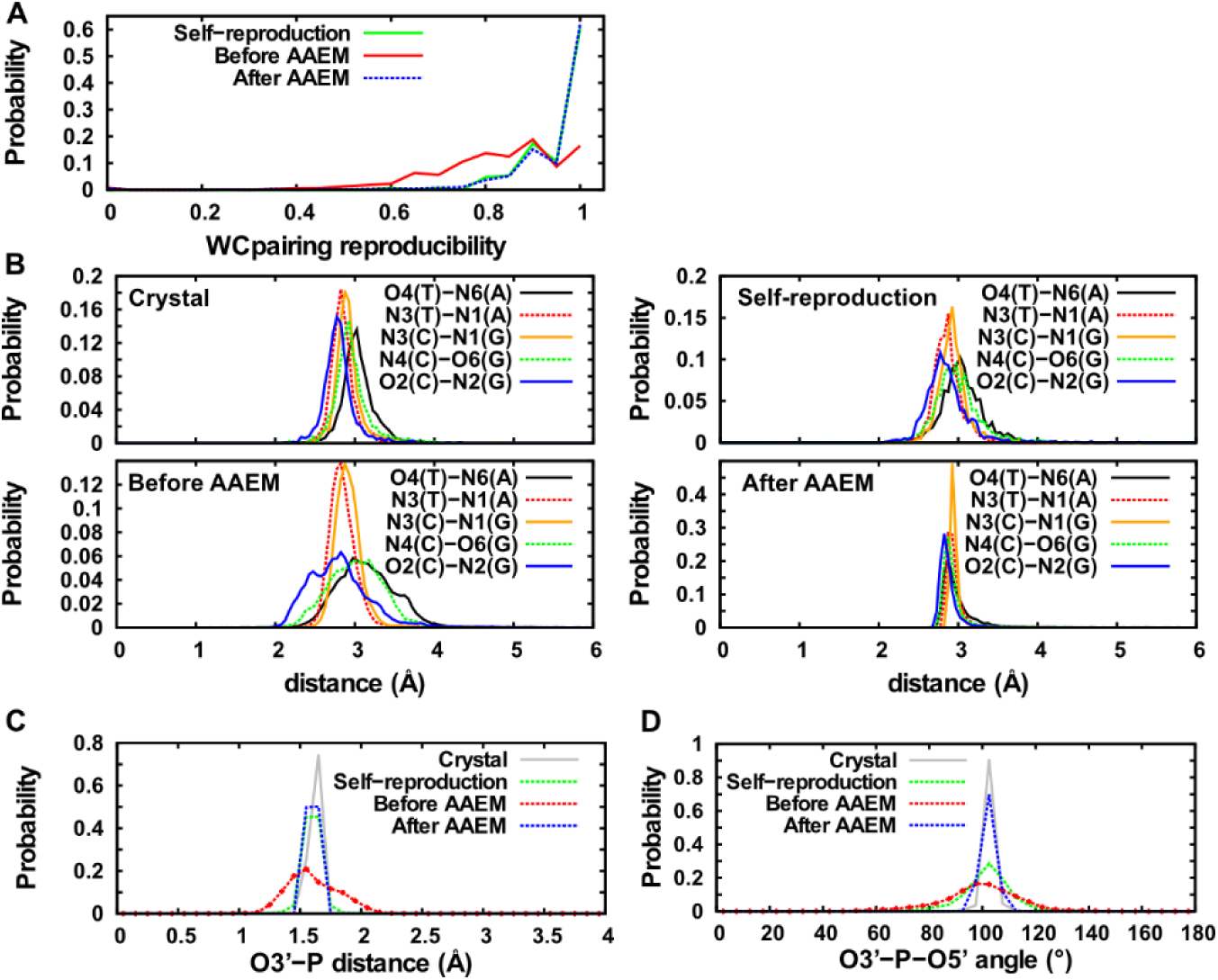
Refinement of local DNA structures by energy minimization with an AA force field for the test-set A. A) The Watson-Crick base pairing reproducibility. B) The distance between donor and acceptor atoms in Watson-Crick pairs. C-D) The distance of O3’-P covalent bond (C) and of O3’-P-O5’ bond angle (D). “Crystal” means crystal structures. “Self-reproduction” means the reconstructed model at self-reproduction test. “Before AAEM” denotes the reconstructed DNA model without the energy minimization and “After AAEM” means the reconstructed model with the subsequent energy minimization. In “After AAEM” case, we used structures which did not stop in following NVT simulation.

We also confirmed these structural improvements are reproducible both by AMBER bsc1 force field and by CHARMM27 force field (Fig S15). While WC pairing reproducibility was slightly low in the CHARMM27 case, the atomistic DNA structure was almost the same. The difference in the force field parameters affects the position of atoms slightly.

In addition to the accurate reproduction of the local DNA structures that we tested so far, the reconstructed model should maintain the DNA bent specific to each DNA-protein complex. Notably, the DNA bent is largely determined by the CG model itself and thus the CG force field is primarily responsible to maintain the correct DNA bend. We examined the reproducibility of the DNA bent in the atomistic model reverse-mapped from CGMD snapshots. In Fig 8A and B, for a TATA binding-protein (PDB ID: 1CDW^69^), we compared the reverse-mapped structure from CGMD snapshots with the original crystal structure. The reconstructed model successfully reproduced the structure of DNA bound to TATA-binding protein. In Fig 8C we calculated the DNA helical axis by Curves+^67,68^: The helical axes in four reconstructed structures (depicted in different colors) reproduced the bents in the DNA helical axis well. In Fig 8D, as another example, we showed the result for the nucleosome (PDB ID: 3UT9^70^), where the reconstructed model reproduced the DNA structure well.

**Figure 8.**
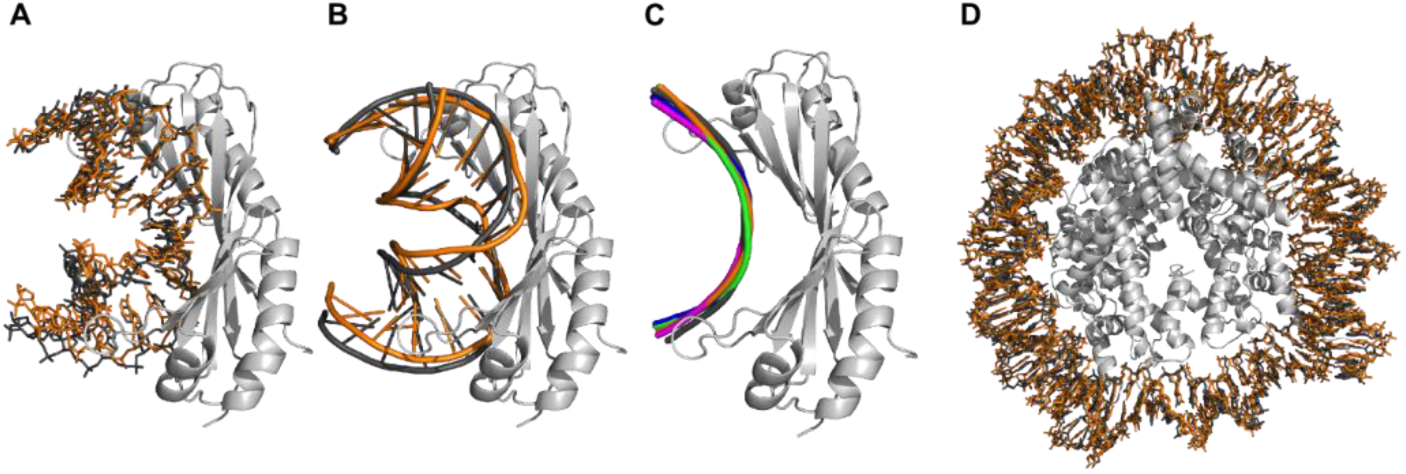
Reproducibility of DNA bending axis of reconstructed model. We compared the crystal structure and reconstructed model after energy minimization at DNA-protein complex. A-C) Comparison of DNA bent by TATA-binding protein (PDB ID 1CDW). A-B) One of reconstructed DNA is superposed on crystal structure and shown in stick model (A) or cartoon model (B) using PyMOL^71^. C) We calculated the DNA helical axes using Curves+. D) Comparison of DNA bound to a nucleosome (PDB ID 3UT9), DNA is shown by stick model of PyMOL. Proteins of crystal structure in light gray, DNA in crystal structure in dark gray. Reconstructed DNA after energy minimization in orange. In C, DNA helical axis of crystal structure in dark gray. The four helical axes of reconstructed models are shown in red, blue, green, and orange, respectively.

### Comparison with other reconstruction methods

We note that suitable reverse-mapping methods strongly depend on the level of coarse-graining. For relatively high-resolution CG models, Rzpiera et al and Wassenaar et al reported general methods for reverse mapping of biomolecular systems^13,14^. In their methods, they allow structural distortion at the first step of atom arrangement. They confirmed that their approach can work well in reconstructing atomic detail from CG protein-lipid model that has a higher resolution than that in this work and in reconstructing a higher-resolution CG lipid model from a lower-resolution CG model. Both methods could reintroduce detailed structure well. Applying the same protocol to our CG model, however, we realized that their pipelines cannot directly be applied to the reconstruction from the Cα based AICG2+ model; proper reconstruction of the sidechain orientation was not realized.

There are several reports discussing reverse-mapping from CG DNA model to atomistic model. Naômé et al, reconstructed atomistic detail from 1 site per nucleotide model^29^. Their method is based on the reverse-mapping by Rzpiera et al. described above^13^. Other groups performed reverse mapping of AA DNA structure from a higher resolution CG model than 3SPN.2C model^15,26,28,45^. In these cases AA structure can be determined nearly uniquely by the CG DNA structure. Our fragment-library-based method can reconstruct a reasonable AA structure from a lower-resolution CG model even before we perform energy minimization with an AA force field. The library-based approach is significantly faster than the MD-based approaches. Therefore our approach can have advantage to reconstruct thousands / millions of structures of flexible molecules (e.g. comparison to small angle X-ray scattering^25^).

We also tried the reverse-mapping for some DNAs using the tool developed by Wassenaar et al, using default parameter except for the correspondence between CG particles and atoms. The program stopped at the energy minimization, or stopped at the process in the MD with LINCS or SETTLE warnings. Probably, more detailed information was necessary to reconstruct from a low resolution CG model. In this sense, our fragment-library based approach is a more efficient procedure.

When one tries to reconstruct AA structure from a low-resolution protein model that does not possess sidechain conformation information, there exist a characteristic difficulty. Once wrong sidechain conformations are introduced in a high-density environment, such as a tightly-packed interface, they cannot easily be corrected. In this context, a general MD-based approach above may have risks. In this study, we attempt to avoid the risks by utilizing sidechain orientations in known 3D structures as much as possible. Indeed, the reproducibility of protein-DNA interaction was significantly superior to the methods that do not use known structure information. Notably, while a direct superposition of known 3D structure has been proposed previously^18^, our method utilizes known structure information only for the sidechains at the interface to DNA. By this way, our approach can incorporate overall conformation changes sampled by CGMD simulations.

## CONCLUSION

In this study, we developed a new reconstruction method of AA structures from CG structural models for protein-DNA complexes. Our primary targets were CG models of protein-DNA complexes represented with one-site-per amino acid in proteins and three-sites-per-nucleotide in DNAs. First we tested our new method for the DNA reconstruction. The method reproduced the tilt of base plane well in most cases and was applicable to non-canonical DNA structures. For snapshots of CGMD simulations, we performed short low-temperature CGMD at 1K before the reconstruction. The Watson-Crick base pairs were well reproduced even though the method does not take into consideration of the hydrogen-bond interaction energy.

Second, we developed the reverse-mapping method from Cα-protein model to atomistic model in which we combined superposition of known atomistic structures with existing modeling tools. By this hybrid method, we successfully reproduced protein-DNA contacts in known 3D structures. The reproducibility of interaction in our hybrid method is equal to or better than the interface obtained after superposition of X-ray crystal structure on CG model. The merit of our approach is that we can treat not only a part where the conformation of sidechains is similar to known structure, but also the other part where the structure is different from experimentally determined structure. This approach should also be applicable to multi-protein systems and other protein-ligand systems.

Finally, we proposed a reverse-mapping method for protein-DNA complexes obtained by prior CGMD simulations. The local structures of atomistic model converged to that observed in X-ray crystal structure. Furthermore, we could perform subsequent constant-temperature AAMD simulations. Therefore, the reverse-mapped structures must have quality usable for further analysis by atomistic MD simulations.

## ASSOCIATED CONTENT

### Supporting Information

The following files are available free of charge.

Details of simulation methods and further analysis of the reconstruction method (PDF)

PDB entries we used for constructing fragment library (Table S1 in SI_Table_S1_S3.xlsx)

PDB entries of dataset we used for testing the reconstruction method (Table S3 in SI_Table_S1_S3.xlsx)

### Notes

The developed program is available at http://www.cafemol.org/download.php.

## AUTHOR INFORMATION

### Funding Sources

The study was supported partly by JSPS KAKENHI grant 25251019(ST), 16KT0054(ST), and 16H01303 (ST), 15H01351(ST) by the MEXT as “Priority Issue on Post-K computer”(ST), and by the RIKEN pioneering Project “Dynamical Structural Biology”(ST).

### Notes

The authors declare no competing financial interest.

## ACKNOWLEDGMENT

We thank Mr. Toru Niina for helping us with debugging our DNA reconstruction program. We also thank Dr. Cheng Tan and Mr. Diego Ugarte La. Torre for useful discussions.

